# R(+) Propranolol decreases lipid accumulation in hemangioma-derived stem cells

**DOI:** 10.1101/2024.07.01.601621

**Authors:** Jerry W.H. Tan, Jill Wylie-Sears, Caroline T. Seebauer, John B. Mulliken, Mathias Francois, Annegret Holm, Joyce Bischoff

## Abstract

**Background:** Infantile hemangioma (IH) is a benign vascular tumor that undergoes an initial rapid growth phase followed by spontaneous involution. A fibrofatty residuum remains in many tumors and often necessitates resection. We recently discovered that R(+) propranolol, the non-β blocker enantiomer, inhibits blood vessel formation of IH patient-derived hemangioma stem cells (HemSC) xenografted in mice. HemSC are multipotent cells with the ability to differentiate into endothelial cells, pericytes, and adipocytes.

**Objectives:** We investigated how R(+) propranolol affects HemSC adipogenic differentiation and lipid accumulation, in vitro and in a preclinical murine model for IH.

**Methods:** We conducted a 10-day adipogenesis assay on 4 IH patient-derived HemSCs. Oil Red O (ORO) staining was used to identify the onset and level of lipid accumulation in HemSC while quantitative real-time polymerase chain reaction was conducted to determine the temporal expression of key factors implicated in adipogenesis. 5-20µM R(+) propranolol treatment was added to HemSC induced to undergo adiogenesis for 4 and 8 days, followed by quantification of lipid-stained areas and transcript levels of key adipogenic factors. We immunostained for lipid droplet-associated protein Perilipin 1 (PLIN1) in HemSC-xenograft sections from mice treated with R(+) propranolol and quantified the area using ImageJ.

**Results:** We found that different patient-derived HemSC exhibit a robust and heterogenous adipogenic capacity when induced for adipogenic differentiation in vitro. Consistently across four IH patient-derived HemSC isolates, R(+) propranolol reduced ORO-stained areas and lipoprotein lipase (LPL) transcript levels in HemSC after 4 and 8 days of adipogenic induction. In contrast, R(+) propranolol had no significant inhibitory effect on transcript levels encoding adipogenic transcription factors. In a pre-clinical HemSC xenograft model, PLIN1-positive area was significantly reduced in xenograft sections from mice treated with R(+) propranolol, signifying reduced lipid accumulation.

**Conclusions:** Our findings suggest a novel regulatory role for the R(+) enantiomer of propranolol in modulating lipid accumulation in HemSC. This highlights a novel role of R(+) propranolol in the involuting phase of IH and a strategy to reduce fibrofatty residua in IH.

**What is already known about this topic?:**

- Propranolol is the mainstay treatment for infantile hemangioma (IH), the most common tumor of infancy, but its use can be associated with concerning β-blocker side effects.
- R(+) propranolol, the enantiomer largely devoid of β-blocker activity, was recently shown to inhibit endothelial differentiation of hemangioma-derived stem cells (HemSC) in vitro and reduce blood vessel formation in a HemSC-derived xenograft murine model of IH.

**What does this study add?:**

- R(+) propranolol inhibits lipid accumulation in HemSC in vitro.
- R(+) propranolol does not affect mRNA transcript levels of key adipogenic transcription factors in differentiating HemSC in vitro.
- R(+) propranolol reduces lipid accumulation in a pre-clinical xenograft murine model of IH.

**What is the translational message?:**

- The R(+) enantiomer of propranolol could be advantageous in terms of reduction in β-adrenergic side effects and fibrofatty tissue formation in the involuting phase of IH.
- Less fibrofatty residua might reduce the need for surgical resection.
- Disfigurement and associated psychosocial impacts might be improved in this young patient cohort.

## INTRODUCTION

Infantile hemangioma (IH) is a common benign vascular tumor that occurs in 5% of infants (1,2). IH arises during early infancy and undergoes rapid vascular growth during the first 6-12 months of infancy (proliferating phase). The involuting phase follows, characterized by spontaneous, slow regression over several years with diminished vascularity and gradual replacement of the tumor with fibrous tissue and islands of fat deposits (3). 10-15% of IHs are considered complex and require treatment.

In 2008, propranolol, a non-selective β-blocker drug, was discovered serendipitously to be an effective therapy for problematic IH (4). A randomized controlled trial showed propranolol induced complete or near complete regression in 60% of IH patients who had received 3 mg/kg/d over 6 months (5). Propranolol treatment has been proposed to accelerate the transition from the proliferating to the involuting phase (6). In long-term follow-up studies, up to 50% of IH patients treated with propranolol or atenolol, a β1-selective blocker, showed fibrofatty tissue residua several years following treatment (7–9).

Propranolol is an equimolar mix of two enantiomers: the S(-) enantiomer showed 1000 times more potent β-adrenergic antagonistic activity than the R(+) enantiomer, which is largely devoid of β-blocker activity at concentrations below 12µM (10). In two prior studies, we demonstrated the R(+) enantiomer of propranolol is sufficient to block endothelial differentiation of IH patient-derived hemangioma stem cells (HemSC) in vitro (11) and IH blood vessel formation in vivo in a pre-clinical xenograft model using HemSC (12). We further showed that R(+) propranolol disrupts the transcriptional activity of sex-determining region Y (SRY) box transcription factor 18 (SOX18), an endothelial-specific transcription factor involved in vascular and lymphatic development and tumor angiogenesis (13–15). In aggregate, these studies suggest a novel β-adrenergic receptor-independent off-target effect of the R(+) propranolol on the transcriptional activities of SOX18 in IH.

The effects of racemic propranolol on fibrofatty residua in IH have been studied in vitro by examining its effects on HemSC adipogenesis (16–18). These studies found that propranolol at doses of 50-100 µM accelerated HemSC adipogenesis initially but caused cell death after 7 days. Possible differential effects of R(+) propranolol on HemSC adipogenesis, however, have not yet been examined.

To address this knowledge gap, we studied the role of R(+) propranolol on HemSC lipid accumulation and adipogenesis. Our findings revealed that R(+) propranolol reduced lipid accumulation in HemSC undergoing adipogenesis for 4 and 8 days.

This was corroborated in vivo in a pre-clinical HemSC xenograft model. The results suggest a novel inhibitory effect of R(+) propranolol on lipid accumulation in IH that may clinically reduce fibrofatty residuum formation.

## MATERIALS AND METHODS

For detailed methods, see Supplementary Methods in Appendix S1.

### IH tissue and cell culture

IH specimens were obtained under an IRB protocol approved by the Committee on Clinical Investigation, Boston Children’s Hospital. IH diagnosis was confirmed by histopathology. Single-cell suspensions were prepared from de-identified proliferating phase IH specimens (Appendix S1). HemSC, hemangioma endothelial cells (HemEC), and human adipose-tissue derived mesenchymal stem cells (AMSC) were isolated and cultured as described (12,19–21).

### Adipogenic differentiation assay

Racemic propranolol and R(+) propranolol (Sigma-Aldrich, USA) were reconstituted at 10mM in phosphate-buffered saline (PBS) to form the stock solution. Rapamycin (LC Laboratories, USA) was reconstituted in DMSO to 10mM to form the stock solution.

On Day -1, HemSC were seeded at 20,000 cells/cm^2^ in fibronectin (0.1 µg/cm^2^)- coated wells on a 24-well plate and allowed to adhere overnight. AMSC were seeded onto 24-well plates without fibronectin. On Day 0, cells were washed with PBS and incubated in adipogenic media (AM), consisting of DMEM-high glucose (Gibco, USA), 10% heat-inactivated fetal bovine serum (FBS; Cytiva, USA), 1x Penicillin-Streptomycin-L-Glutamine (GPS; Corning, USA), 1µM dexamethasone (Sigma-Aldrich, USA), 0.5mM 3-isobutyl-1-methylxanthine (IBMX; Sigma-Aldrich, USA), 5µg/ml insulin (SAFC, USA) and 60µM indomethacin (Sigma-Aldrich, USA). AM without dexamethasone, IBMX, insulin, and indomethacin served as the control, termed “Control Media” in adipogenesis assays. Media were replaced every 48 hours.

### Oil Red O-staining

To prepare the Oil Red O (ORO; Sigma-Aldrich, USA) working solution, a 3:2 ratio of 0.5% (w/v) ORO in isopropanol (Fisher Scientific, USA): double distilled H_2_O (ddH_2_O) was freshly prepared and filtered to remove precipitates.

For ORO staining of lipid droplets, 12mm coverslips (Electron Microscopy Sciences, USA) were coated with fibronectin (0.1 µg/cm^2^) in 24-well plates before seeding of HemSC. Cells were fixed in 4% paraformaldehyde (Electron Microscopy Sciences, USA) for 15 minutes at RT, washed in PBS, and incubated in 60% isopropanol for 3 minutes. Cells were stained with ORO for 10 minutes at RT, and washed with 60% isopropanol for 10 minutes to remove unbound ORO dye. Cells were washed twice in ddH_2_O, mounted onto slides, and imaged within 24-48 hours to avoid precipitation.

### Quantitative real-time polymerase chain reaction (qRT-PCR)

Total RNA was extracted from cells using the RNeasy micro kit (Qiagen, Netherlands). cDNA was synthesized with iScript cDNA Synthesis Kit (Bio-Rad, USA) as per manufacturer’s protocol. qRT-PCR was performed using the KAPA SYBR FAST ABI Prism 2x qPCR Master Mix (Roche, Switzerland) and a QuantStudio 6 Flex Real-Time PCR system (Thermo Fisher Scientific, USA). Primer pairs with an amplification efficiency of 90-110% were selected (Appendix S1). ATP Synthase F1 Subunit β (ATP5B) was used as a housekeeping gene. Fold changes in gene expression were determined using the ΔΔCt method, normalized against ATP5B Ct values. Each amplification reaction was performed in triplicate.

### In vivo murine model for IH vasculogenesis

The IH xenograft experiment was conducted as described by Seebauer et al. (2022). The murine xenograft tissue slides were used for immunofluorescent staining with anti-human perilipin-1 (PLIN1) as a surrogate for lipid content. Full details are provided in Appendix S1.

### Immunohistochemistry

Formalin-fixed paraffin-embedded (FFPE) tissue sections (5μm) of Matrigel implants were deparaffinized and immersed in a Citrate-EDTA antigen retrieval buffer (pH 6) for 20 minutes at 95°C–99°C. Sections were blocked for 30 minutes in 10% donkey serum and incubated with primary and secondary antibodies (Appendix S1).

### Imaging and Quantification

For imaging of ORO-stained slides, brightfield color images were taken through a 20x objective lens using the Zeiss Axio Imager M1 microscope (Carl Zeiss Microscopy, USA). ORO-stained areas were determined from 5 fields/replicate with 2 replicates per treatment condition. Each field was 706.56 µm x 518.88 µm = 0.366 mm^2^.

For imaging of PLIN1-stained sections, confocal images were acquired by an LSM 880 confocal microscope (Carl Zeiss Microscopy, USA) using a 20x objective lens. PLIN1-stained area and total nuclei count were determined from 5 fields/section, 1 section/implant, and 2 implants/mouse. Each field was 425.1 µm x 425.1 µm = 0.18071 mm^2^.

To minimize bias associated with imaging, blinded image analysis using anonymous labeling was conducted when imaging representative fields. PLIN1-stained area, total nuclei count, and ORO-stained area were analyzed using FIJI ImageJ software (NIH, USA).

### Statistical Analysis

Results are shown as mean ± SD. Differences considered significant at P<0.05. One-way ANOVA was used to analyze for differences between multiple groups while Student’s two-tailed test was used for comparisons between two groups. When data do not follow Gaussian distribution, non-parametric Kruskal-Wallis test or Mann-Whitney test were used instead. GraphPad Prism was used for analysis. FIJI ImageJ was used for densitometric analysis of western blot data. qRT-PCR data from HemSC adipogenic differentiation test was transformed as detailed by Willems et al. (2008).

## RESULTS

### Different IH patient-derived HemSC display heterogeneous potential for adipogenic differentiation in vitro

HemSC have been shown to undergo adipogenic differentiation in vitro, demonstrated by lipid accumulation (19), increased lipoprotein lipase (LPL), and CCAAT-enhancer binding protein alpha (C/EBPα) transcript levels (16,22). To better understand HemSC adipogenic capability, we conducted a 10-day time course to identify the onset and level of lipid accumulation and expression of transcription factors implicated in adipogenesis. HemSC isolated from 4 different IH specimens were differentiated in adipogenic media (23). In parallel, HemSC were incubated in control media. Cells were stained with ORO on Day 0, 2, 4, 6, 8 and 10. Semi-quantitative image analysis was conducted to assess the intracellular neutral lipid droplets. AMSC were used as a positive control (20).

HemSC isolated from 4 different IH showed heterogeneity in the levels of ORO-positive areas during the 10 days of induction (Figure 1a,b; Figure S1). As early as Day 4 of adipogenic induction, visible lipid droplets were observed in all HemSC (Figure 1a).

**Figure 1:**
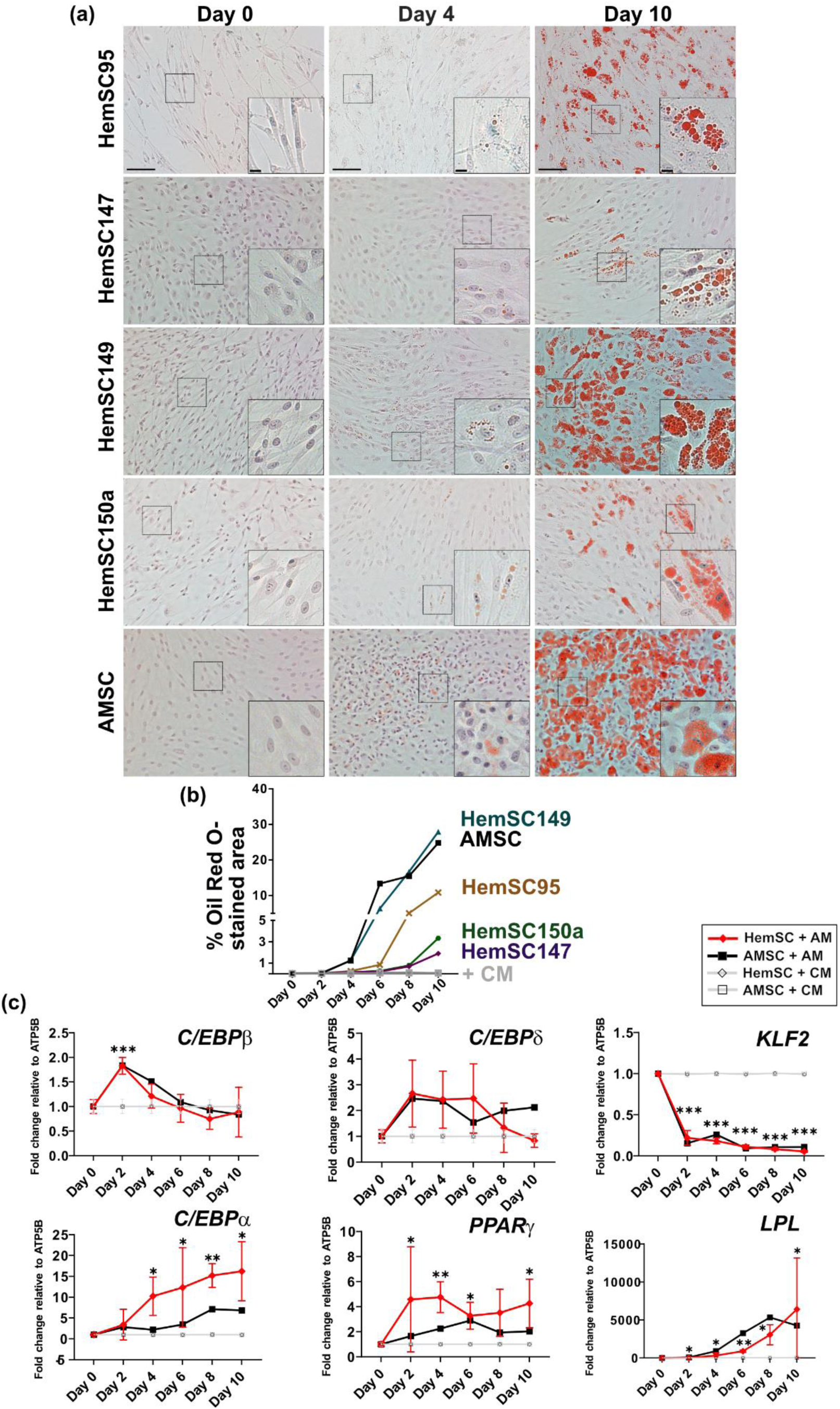
Hemangioma stem cells (HemSC) display heterogeneous potential for adipogenic differentiation in vitro (a) Representative Oil Red O (ORO) images of HemSC (n=4) and human adipose tissue-derived mesenchymal stem cells (AMSC) (n=1) at Day 0, 4, and 10 of a 10-day adipogenic differentiation in vitro assay. Scale bar = 100µm. Magnified images from region of interest (denoted by black dotted box) shown (bottom right), scale bar = 15µm. (b) Quantification of ORO-stained area as a percentage of total image area shown in (a). ORO-stained area of cells grown in Control Media (CM) (not shown) and Adipogenic Media (AM) (shown in Figure S1). (c) Gene expression of CCAAT-enhancer-binding protein beta (C/EBPβ), C/EBPδ, C/EBPα, peroxisome proliferator-activated receptor gamma (PPARγ), Kruppel-like factor 2 (KLF2), and lipoprotein lipase (LPL) determined by quantitative real time-polymerase chain reaction. Data presented as mean ± SD, n=4 for HemSC and n=1 for AMSC. *P<0.05, **P<0.01, ***P<0.001 compared to cells grown in CM for the corresponding duration.

In parallel, qRT-PCR was conducted to quantify transcript levels of key adipogenesis-related transcription factors in the CCAAT-enhancer-binding protein (C/EBP) family (C/EBPβ, C/EBPδ, C/EBPα) as well as peroxisome proliferator-activated receptor-γ (PPARγ) (24). LPL levels were determined to investigate expression levels of genes regulated by the PPARγ and C/EBP family of transcription factors. The anti-inflammatory endothelial transcription factor Kruppel-like factor 2 (KLF2) was recently discovered to be a regulator of HemSC differentiation (25) and is a negative regulator of adipogenesis (26). For this reason, KLF2 was also quantified by qRT-PCR.

Early-expressing adipogenic factors C/EBPβ and C/EBPδ were upregulated after 2 days of induction (Figure 1c). C/EBPβ levels declined beyond Day 2 while C/EBPδ levels were sustained until Day 6 and decreased from Day 6 to Day 10. Terminal transcription factors C/EBPα and PPARγ were significantly upregulated from Day 4 and remained elevated at Day 10. In contrast, KLF2 levels decreased significantly after 2 days of adipogenic induction and remained low thereafter (Figure 1c). LPL levels increased exponentially from Day 4 to Day 8 of induction onwards in HemSC. In summary, by Day 8, HemSC displayed extensive cytoplasmic lipid droplet formation (Figure S1), showed upregulated C/EBPα, PPARγ, and LPL, and downregulated KLF2. When compared to AMSC, all HemSC except for HemSC149 showed a modest lipid accumulation profile.

### R(+) propranolol and racemic propranolol reduce lipid accumulation in differentiating HemSCs in vitro

The time course study suggested that differentiating HemSC at Day 4 were transitioning to an adipocyte phenotype based on appearance of lipid droplets and increased C/EBPα and PPARγ expression. HemSC at Day 8 had formed extensive lipid accumulation consistent with further maturation to an adipocyte phenotype. Therefore we chose Day 4 and Day 8 to study the effects of R(+) propranolol and racemic propranolol on adipogenic differentiation of HemSC in vitro.

We cultured HemSC in adipogenic media for 4 and 8 days with 0-20µM R(+) propranolol or racemic propranolol. The range was selected based on prior studies in which 10μM propranolol blocked norepinephrine-induced dilation of human coronary arterial segments (27). Furthermore, 5μM propranolol was sufficient to block HemSC to endothelial differentiation in vitro (11). The mammalian target of rapamycin (mTOR) inhibitor, rapamycin at 20ng/ml was used as a positive control as it inhibits lipid accumulation and adipogenic differentiation of HemSC (22).

On Day 4, ORO staining was reduced in differentiating HemSC cultured in R(+) propranolol or racemic propranolol compared to PBS-treated controls (Figure 2a; Figure S2). Quantification showed significant reductions in the percentage of ORO-stained area in differentiating HemSC cultured in R(+) propranolol and racemic propranolol at all doses (5-20µM) as compared to PBS-treated controls (Figure 2b and c). On Day 8, the inhibitory effects of R(+) propranolol and racemic propranolol on ORO-stained areas were modest (Figure 2d). Quantification of ORO-stained areas showed a significant reduction at 10 and 20µM R(+) propranolol and 20µM racemic propranolol compared to PBS-treated controls (Figure 2e and 2f). The positive control rapamycin was more effective than either racemic or R(+) propranolol at inhibiting HemSC lipid accumulation after both 4 days and 8 days of adipogenesis (Figure S2). In summary, racemic and R(+) propranolol inhibited lipid accumulation of HemSC undergoing adipogenesis in vitro on both Day 4 and Day 8.

**Figure 2:**
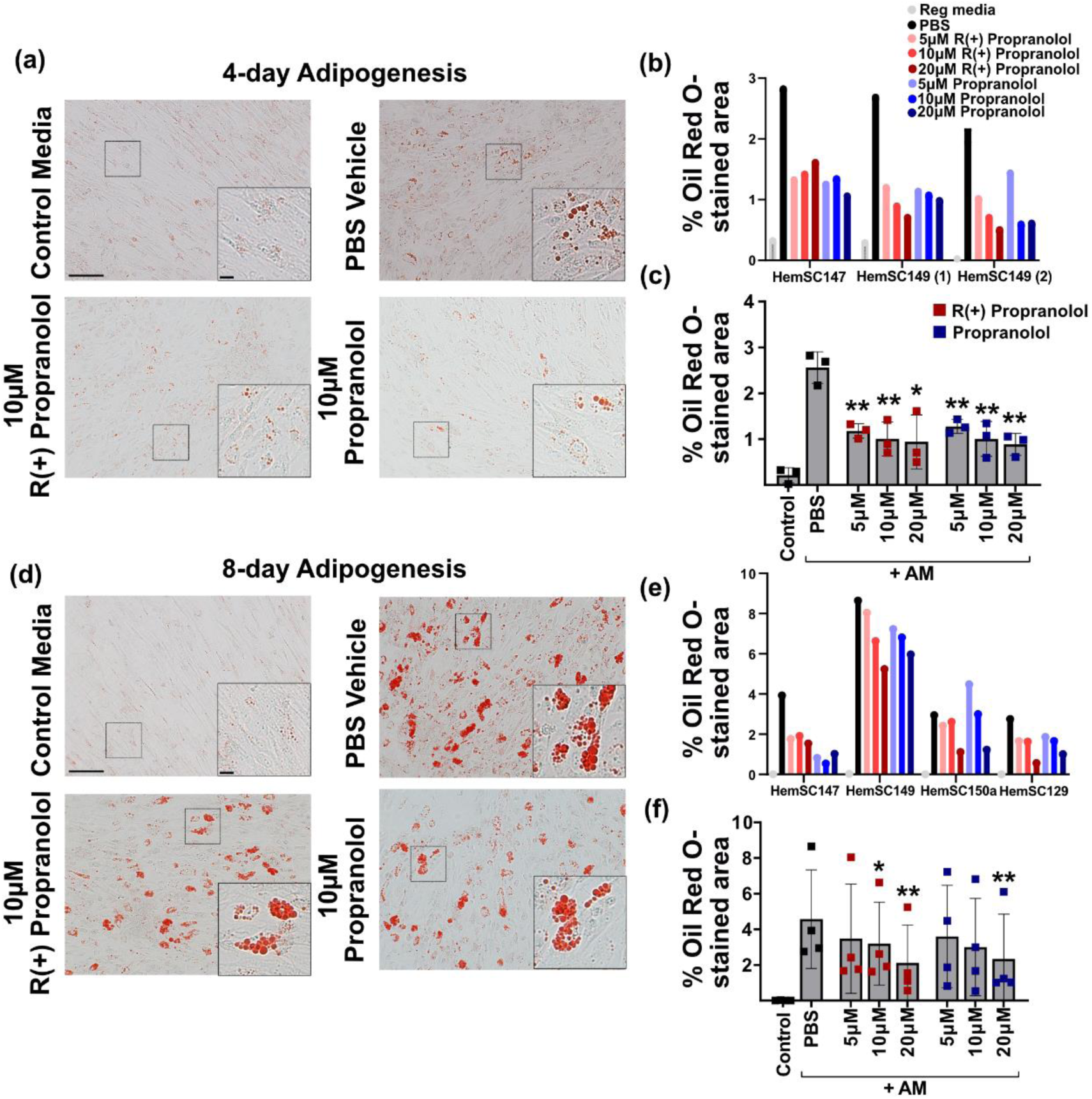
R(+) propranolol and racemic propranolol decrease lipid accumulation in Hemangioma stem cells (HemSC) undergoing adipogenesis in vitro. HemSC grown in adipogenic media (AM) for 4 days (a-c) and 8 days (d-f) in 0 – 20µM R(+) propranolol or racemic propranolol. (a) Representative Oil Red O (ORO) images (Shown: HemSC149 (2) from (b)) of HemSC undergoing 4 days of adipogenesis with phosphate-buffered saline (PBS) vehicle, 10µM R(+) propranolol, and 10µM racemic propranolol. HemSC grown in control media (CM) were used as a negative control. Scale bar = 100µm. Magnified areas from region of interest (denoted by black dotted box) shown (bottom right), scale bar = 15µm. (b) Quantification of % ORO-stained area for each HemSC line (147 and 149) over n=3 independent experiments. (c) Average ORO-stained area (% total) from (b). Data presented as mean ± SD, n= 3. *P<0.05, **P<0.01 compared to cells grown in AM with PBS as vehicle control. (d) Representative ORO images of HemSC (shown: HemSC149 from (e)) undergoing 8 days of adipogenesis, scale bar = 100 µm. Magnified areas at lower right, scale bar = 15 µm. HemSC grown in AM for 8 days in 5–20µM R(+) propranolol or racemic propranolol. (e) Quantification of % ORO-stained area in each HemSC lines (147, 149, 150a, 129) over n=4 independent experiments. (f) Average ORO-stained area (% total) from (e). Data presented as mean ± SD, n=4. *P<0.05, **P<0.01 compared to cells grown in PBS vehicle in AM.

### Reduction in lipid accumulation by R(+) propranolol is not associated with reduced transcript levels for adipogenic transcription factors

To determine if the reduced lipid accumulation in R(+) propranolol-treated differentiating HemSC is due to a disruption in adipogenic differentiation, we assayed the transcript levels of the key adipogenic transcription factors C/EBPβ, C/EBPδ, C/EBPα, and PPARγ, and the endothelial transcription factor KLF2 in drug-treated cells. Strikingly, there were no significant reductions in mRNA levels of these transcription factors on Day 4 or Day 8 of adipogenesis (Figure 3a). On Day 4, C/EBPβ, C/EBPδ, and C/EBPα expression levels remained largely unchanged while PPARγ expression showed a significant but modest increase in expression in differentiating HemSC treated with 5-20µM R(+) propranolol or 5-10µM racemic propranolol (Figure 3a). Anti-adipogenic transcription factor KLF2 also remained unchanged in R(+) propranolol and racemic propranolol-treated differentiating HemSC. The adipogenic marker LPL was reduced in HemSC treated with 10-20µM R(+) propranolol or 20µM racemic propranolol (Figure 3a). Treatment with rapamycin for 4 days had no effect on levels of early adipogenic markers C/EBPβ and CEBPδ, yet caused a reduction in terminal markers C/EBPα, and LPL, as reported previously (22), and increased KLF2 levels (Figure S2).

**Figure 3:**
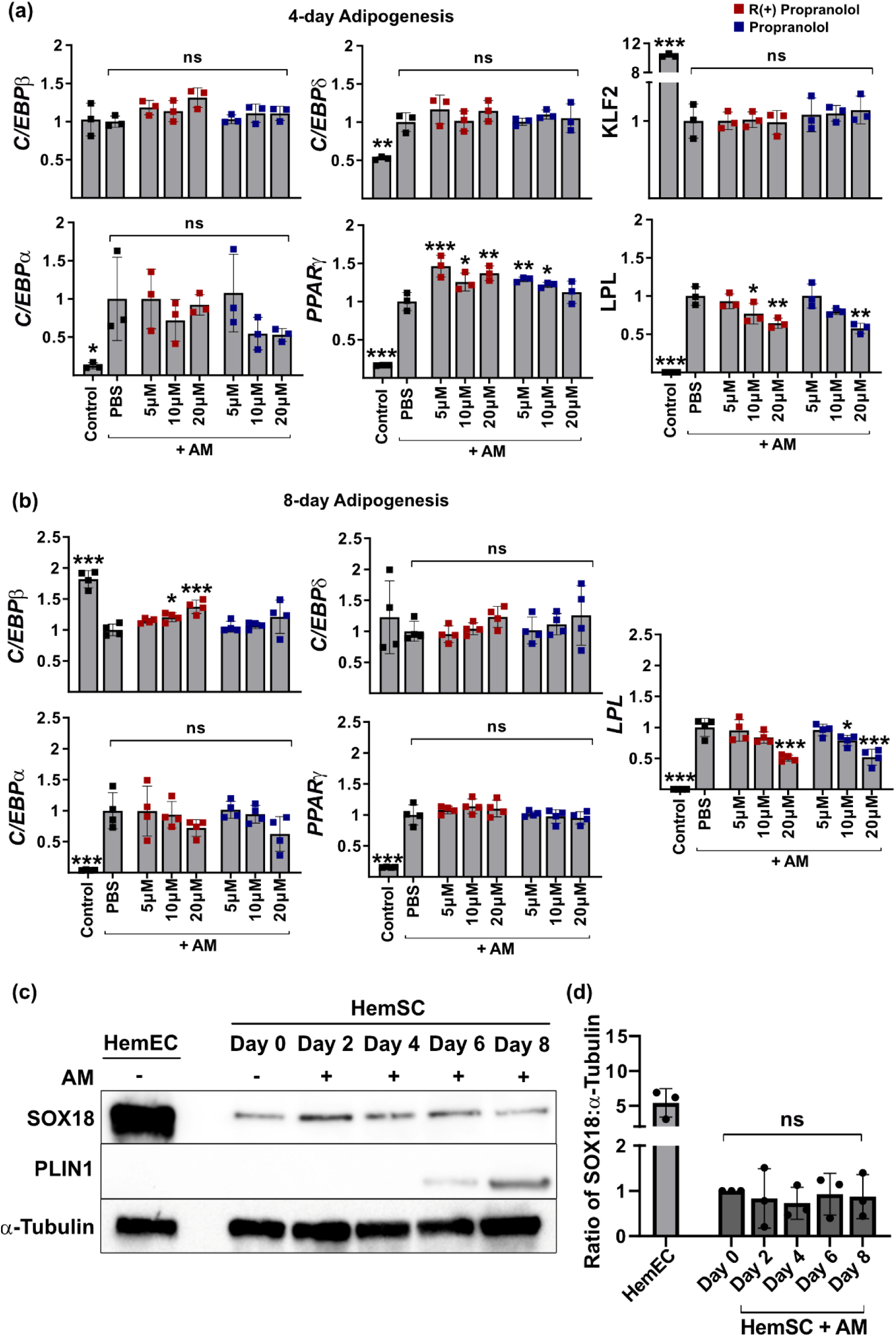
R(+) propranolol did not reduce mRNA levels of key adipogenic transcription factors in HemSC undergoing adipogenic differentiation. (a) Total RNA extracted from 4 day-cultured cells from (Figure 2a) and gene expression of adipogenesis-related genes normalized against that of cells grown in adipogenic media (AM) with phosphate-buffered saline (PBS) vehicle. Gene expression of CCAAT-enhancer-binding protein beta (C/EBPβ), C/EBPδ, C/EBPα, peroxisome proliferator-activated receptor gamma (PPARγ), Kruppel-like factor 2 (KLF2), and lipoprotein lipase (LPL) determined by quantitative real time-polymerase chain reaction. Data are presented as mean ± SD, n=3 (b) Total RNA extracted from 8 day-cultured cells from (Figure 3d) and gene expression of adipogenesis genes normalized against that of cells grown in AM with PBS vehicle. Data presented as mean ± SD, n=4. *P<0.05, **P<0.01, ***P<0.001 compared to cells grown in PBS. (c) Western Blot for Sry-box transcription factor 18 (SOX18) and Perilipin-1 (PLIN1) in HemSC149 undergoing adipogenic differentiation over 8 days. Protein lysates from hemangioma endothelial cell (HemEC) analyzed as positive control for SOX18. (d) ImageJ quantification of SOX18 protein levels in n=3 HemSC (149,171,150a) normalized to α-tubulin. Data presented as mean ± SD, n=3. Full blots shown in Supplementary Figure S3.

On Day 8, mRNA levels of C/EBPδ, C/EBPα, and PPARγ were unchanged after treatment with R(+) propranolol and racemic propranolol compared to PBS vehicle-treated HemSC induced to undergo adipogenesis. C/EBPβ was significantly but weakly increased by 10 and 20μM R(+) propranolol compared to PBS-treated HemSC induced to undergo adipogenesis. On Day 8, LPL was reduced when HemSC undergoing adipogenesis were treated with 20µM R(+) propranolol or 10-20µM racemic propranolol (Figure 3b). Rapamycin treatment for 8 days resulted in a significant upregulation of early marker C/EBPδ and significant downregulation of terminal markers C/EBPα, PPARγ, and LPL (Figure S2). These results demonstrate that the R(+) propranolol and racemic propranolol-induced inhibition of lipid accumulation in differentiating HemSC does not coincide with a reduction in transcript levels encoding adipogenic transcription factors.

To ascertain if the inhibitory effect of R(+) propranolol and racemic propranolol on lipid accumulation is HemSC-specific, we conducted an 8-day in vitro adipogenesis assay on AMSC (Figure S4). This experiment showed a slight reduction in the percentage of ORO-stained area in differentiating AMSC treated with 20µM R(+) propranolol, similar to that observed in 20µM R(+) propranolol-treated differentiating HemSC (Figure 2e and 2f). Conversely, lipid accumulation was not reduced in racemic propranolol-treated AMSC, and dose-dependent changes were not observed (Figure S4). Expression levels of transcription factor C/EBPβ, C/EBPδ, and PPARγ were unchanged whereas C/EBPα and LPL levels fell with 20µM R(+) and racemic propranolol treatment. This is similar to what was observed in differentiating HemSC-treated with 20µM R(+) and racemic propranolol for 8 days (Figure 3b). In summary, R(+) propranolol demonstrates a similar inhibitory effect on lipid accumulation in both HemSC and AMSC and similarly does not appear to affect mRNA levels of transcription factors involved in adipogenesis.

At 20µM, R(+) propranolol was found to inhibit the dimerization of SOX18 transcription factor, thereby inhibiting SOX18 transcriptional activity in vitro and in vivo (11,12). To determine if SOX18 is present in HemSC over 8 days of adipogenic differentiation and thus a potential target, HemSC cell lysates isolated on Day 0, 2, 4, 6, and 8 of adipogenic differentiation were analyzed by Western Blot. The lipid droplet/adipocyte marker PLIN1 was analyzed in parallel (Figure S3).

SOX18 protein was detected in HemSC (n=3) at Day 0 and remained expressed at a basal level throughout 8 days of adipogenesis (Figure 3c and 3d). The SOX18 levels seen in HemSC were low, as expected, compared to levels detected in HemEC (Figure 3c and 3d). PLIN1 was detected on Day 6 and Day 8, consistent with lipid accumulation (Supplemental Figure S3). The protein expression of SOX18 by HemSC undergoing adipogenic differentiation suggests that the reduction in lipid accumulation by R(+) propranolol could potentially be mediated by the inhibition of SOX18 transcriptional activity.

### R(+) propranolol decreases lipid accumulation in a pre-clinical IH xenograft murine model

Seebauer et al. (2022) showed that R(+) propranolol inhibits blood vessel formation in a murine xenograft model of IH (Figure 4a). Cells with lipid droplets were observed in Hematoxylin and Eosin (H&E) stained sections of HemSC xenografts removed on Day 8 in this study. To determine if R(+) propranolol affected lipid accumulation in vivo, we immunostained the xenograft sections with anti-human PLIN1. PLIN1 coats the periphery of lipid droplets and thereby serves as a proxy for intracellular neutral lipid droplets (28). To ensure no cross-reactivity with murine PLIN1, we immunostained mouse adipose tissue sections with the anti-human PLIN1 to determine the dilution of anti-human PLIN1 antibody that would not show cross-reactivity with murine PLIN1 (Figure S5).

**Figure 4:**
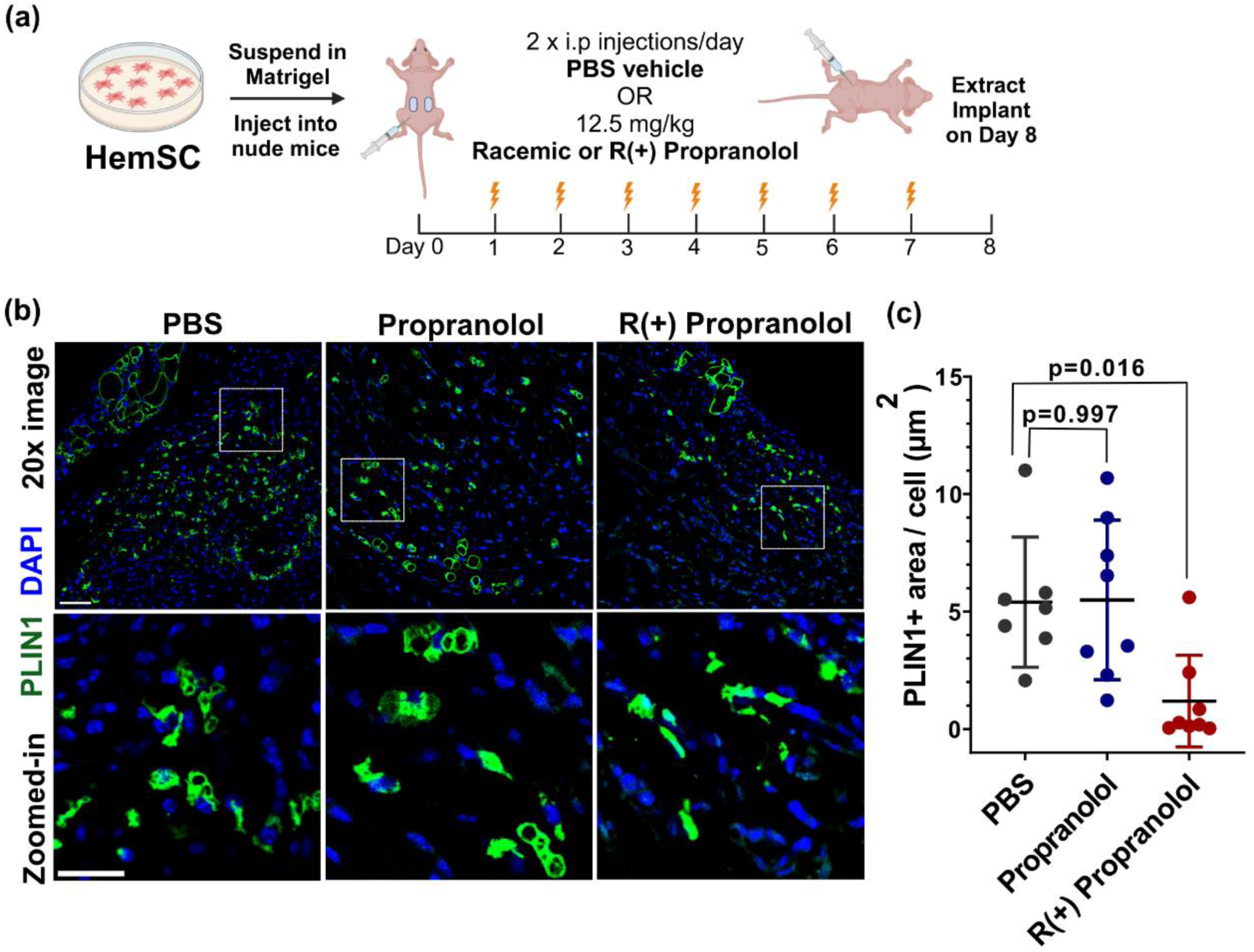
R(+) propranolol decreases lipid accumulation in pre-clinical infantile hemangioma (IH) xenograft model. (a) Schematic of sequential steps involved in IH xenograft model. Hemangioma stem cells (HemSC) pre-treated with phosphate-buffered saline (PBS) or 10 µM of propranolol or R(+) propranolol, suspended in Matrigel, and injected subcutaneously into nude mice, with 2 implants per mouse (n = 8 implants for each group). Mice treated with 12.5 mg/kg drug versus PBS vehicle twice a day for 7 days. The HemSC xenograft implants harvested on Day 8. Figure created with BioRender.com. (b) Immunofluorescent staining of formalin-fixed paraffin-embedded sections for anti-human perilipin-1 (PLIN1). Top: Representative 20x images. Scale bar = 50µm. Bottom: Magnified images at region of interest shown by white box. Scale bar = 25µm. (c) Quantification of PLIN1-stained area per cell (µm^2^) of implants in (b). Data presented as mean ± SD, n=7-8.

We noticed numerous PLIN1-positive cells in the HemSC implants of PBS-treated control mice (Figure 4b). Human PLIN1-positive area per cell was significantly reduced in the implants from 12.5 mg/kg R(+) propranolol-treated mice (PBS: 5.402 µm^2^ per cell vs R(+) propranolol: 1.191 µm^2^ per cell, p-value = 0.016) but not in implants from mice treated with 12.5 mg/kg racemic propranolol (Propranolol: 5.497 µm^2^ per cell, p-value = 0.997) (Figure 4c). A trend towards reduced PLIN1 was seen in implants from mice treated with 5 mg/kg R(+) propranolol, indicating dose response (Figure S6). These data suggest that the R(+) enantiomer of propranolol, but not racemic propranolol, decreases lipid accumulation dose-dependently in our pre-clinical xenograft model of IH.

## DISCUSSION

Prior studies have elucidated the molecular pathways defining the in vitro differentiation of HemSC into endothelial cells and pericytes (29–31), but not adipocytes, the primary cellular component in involuting IH. Furthermore, the effect of the R(+) enantiomer of propranolol on IH involution and fibrofatty tissue formation has not been studied; here we address this knowledge gap.

We demonstrate that R(+) propranolol decreases lipid accumulation in IH patient-derived HemSC in vitro and in vivo. Notably, the R(+) propranolol effect on lipid accumulation in HemSC induced to undergo adipogenesis did not coincide with a decrease in transcript levels of key adipogenic transcription factors, suggesting that R(+) propranolol could directly modulate pathways regulating lipid synthesis or lipolysis rather than adipogenic differentiation *per se*.

Racemic propranolol, an equimolar mix of the R(+) and S(-) enantiomers, also reduces lipid accumulation in HemSC undergoing adipogenic induction in vitro. This differs from previous studies that report propranolol enhanced lipid accumulation after 4 days of adipogenic induction and caused dysregulated HemSC adipogenesis (16–18). Of note, these in vitro studies used propranolol doses in the range of 50– 100µM, which could cause toxicity. In our study, 5–20µM racemic propranolol resulted in a similar reduction in lipid accumulation in differentiating HemSC as its non-β-blocker R(+) enantiomer, which we attribute to the presence of the R(+) enantiomer in racemic propranolol.

Intriguingly, racemic propranolol had no effect on the extent of PLIN1 staining in the HemSC xenografts after 7 days in vivo. In contrast, R(+) propranolol caused a significant reduction in PLIN1 staining. Detailed cellular and molecular studies are needed to understand the competing activities of SOX18 disruption and β-adrenergic receptor blockade in mice treated with racemic propranolol.

Of note, we observed heterogeneous lipid accumulation potential among HemSC isolated from different patient IH specimens, which may be explained by variations in tumor stage at time of resection, location, and gender. Despite these inherent differences, HemSC are comparable to AMSC over a 10-day time course of adipogenic induction. We posit that the adipogenic differentiation process consists of an initial commitment to the adipogenic lineage in the first 2 days of induction before reaching adipocyte maturation after 8 days of induction.

Further insights into the mechanism of propranolol on lipid metabolism can be gleaned from prior studies on pre-adipocytes. For example, β-adrenergic receptor activation during metabolic stress has well-known effects in stimulating lipolysis to release intracellular free fatty acids. In 3T3-L1 adipocytes, the β-adrenergic receptor agonist isoproterenol stimulates cAMP production and lipolysis, while rescue with propranolol inhibits this effect. Propranolol treatment without isoproterenol, however, did not affect basal lipolysis rates (28), suggesting that the β-adrenergic antagonism may not be relevant to lipid accumulation in HemSC. Propranolol was also found to disrupt triglyceride synthesis by inhibiting Lipin-1, an enzyme involved in catalyzing the conversion of phosphatidic acid (PA) to diacylglycerol in the penultimate step of triglyceride synthesis (32). Knockdown of Lipin-1 leads to PA accumulation that inhibits lipid accumulation in 3T3-L1 pre-adipocytes during early differentiation but not during terminal differentiation of mature adipocytes (32).

Interestingly, we observed a greater inhibition of lipid accumulation on Day 4 compared to Day 8 in HemSC undergoing adipogenesis with either R(+) or racemic propranolol. This suggests a time-sensitive role for propranolol in regulating lipogenesis at specific developmental stages of HemSC adipogenesis.

Our recent findings using bulk RNA-seq identified differentially expressed genes in HemSC undergoing endothelial differentiation treated ± R(+) propranolol, including downregulation of genes involved in cholesterol biosynthesis such as the rate-limiting enzyme 3-hydroxy-3-methylglutaryl coenzyme A reductase (Holm et al., bioRxiv; doi.org/10.1101/2024.01.29.577829). The mevalonate pathway, responsible for cholesterol biosynthesis, was reported to be critical in adipocyte survival (33,34), which could provide early clues to the mechanistic role of R(+) propranolol and SOX18 in adipocyte biology. However, direct evidence for SOX18 expression in adipocytes is lacking. A limitation of our data is that Western blot detection of SOX18 in the HemSC cultures undergoing adipogenic differentiation does not distinguish between SOX18 in non-differentiated versus differentiated HemSC.

Propranolol is the mainstay of treatment for complex IH. While propranolol is effective in the majority of IH patients, safety concerns regarding short and long term side-effects attributed to its β-blocker activity in this vulnerable pediatric patient cohort are emerging. R(+) propranolol was recently shown to inhibit IH blood vessel formation in vivo during the proliferating phase of IH. These findings together with results presented here provide the groundwork to improve upon the success of propranolol therapy for IH.

In conclusion, our findings provide insights into a novel role of the R(+) enantiomer of propranolol in inhibiting lipid accumulation in HemSC in vitro and in vivo. We hypothesize that these processes may be regulated initially through transcriptional inhibition of SOX18 by R(+) propranolol, however, unknown off-target activities may be in play. Experiments examining the potential role of SOX18 in the lipid metabolism of differentiating HemSC will illuminate our understanding of the underlying molecular basis. Finally, our findings propose important clinical impacts and R(+) propranolol may offer two therapeutic benefits: Diminished β-adrenergic side effects, and possibly less fibrofatty tissue accumulation in the involuting phase of IH.

## Acknowledgements

We thank Sana Nasim for technical guidance, Andrew Kuo for technical guidance and critical reading of the manuscript, and Aram Ghalali and Hiroko Kishikawa for technical expertise. We thank Allen Chilun Luo and Juan Melero-Martin for gifting AMSC and providing technical expertise and Dr. Roopali Roy, for providing murine fat tissue.

## Data Availability Statement

No datasets were generated or used in this study

