## Supplemental Figures for "R(+) Propranolol decreases lipid accumulation in hemangioma-derived stem cells"

**SUPPLEMENTARY TABLES**

**Appendix S1**

| Sample ID | Gender | Age at Resection (months) | Figure |
|---|---|---|---|
| Proliferating IH (<1 year) | | | |
| 150a | M | 3 | 1,2,3,4 |
| 147 | M | 3 | 1,2,3 |
| 149 | F | 7 | 1,2,3 |
| 95 | F | 8 | 1,2,3 |
| 129 | F | 9 | 2,3 |
| 171 | F | 4 | 3 |

**Table S1: Infantile Hemangioma (IH) Patient Characteristics**

| Gene | Forward | Reverse |
|---|---|---|
| ATP5B | ccactaccaagaagggatctatca | gggcagggtcagtcaagtc |
| CEBPβ | ggtttcgaagttgatgcaatcg | caacaagcccgtaggaacat |
| CEBPδ | actcagcaacgacccatacc | cgctcctatgtcccaagaaa |
| CEBPα | tggacaagaacagcaacgag | ttgtcactggtcagctccag |
| PPARγ | gctggcctccttgatgaata | ttgggctccataaagtcacc |
| LPL | gtcagagccaaaagaagcagc | atgggtttcactctcagtccc |
| KLF2 | *ccacgatcctccttgacgag* | *ccgcagacagtacaaattaaggc* |

**Table S2: Quantitative real time-polymerase chain reaction (qRT-PCR) primers.**

### SUPPLEMENTARY FIGURES

### Appendix S1

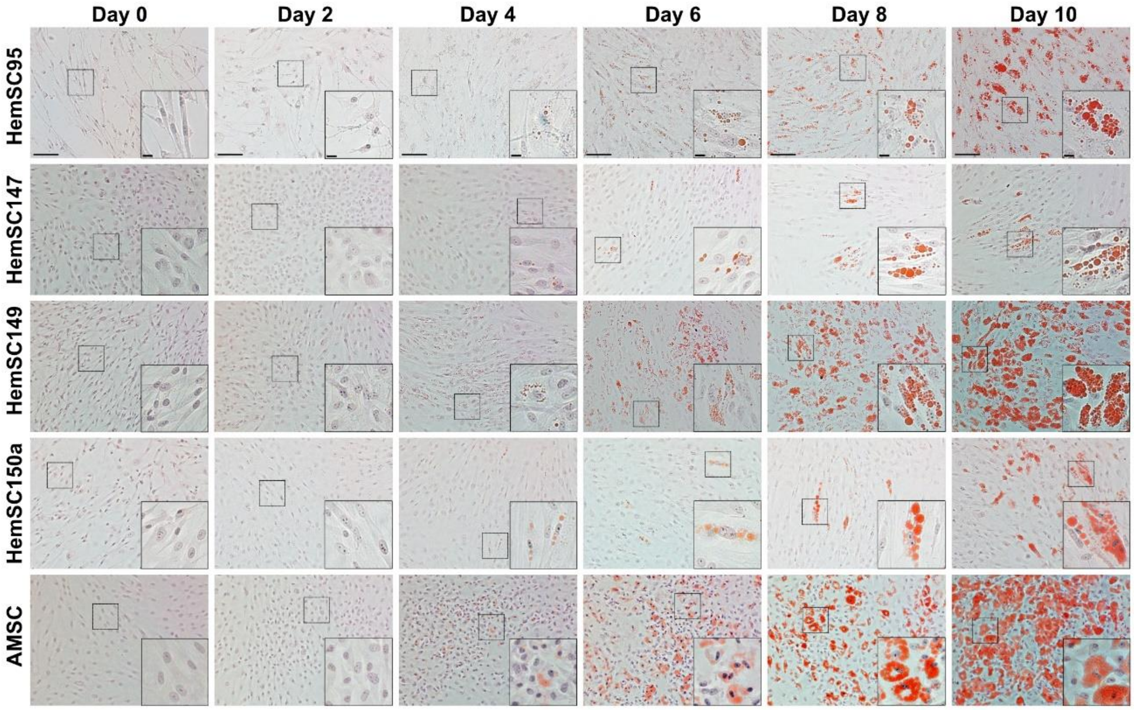


**Figure S1: Time course characterization of Hemangioma-derived stem cell (HemSC) undergoing adipogenesis in vitro.** Representative Oil Red O images of HemSC (N = 4) and human white adipose tissue-derived mesenchymal stem cell (AMSC) (N = 1) at Day 0, 2, 4, 6, 8, and 10 of adipogenic differentiation in vitro assay. Each HemSC isolated from the hemangioma tissue from a unique patient, deidentified and numbered as 95, 147, 149, 150a. Scale bar = 100 μm. Magnified images from regions of interest denoted by black dotted box. Scale bar = 15 μm.

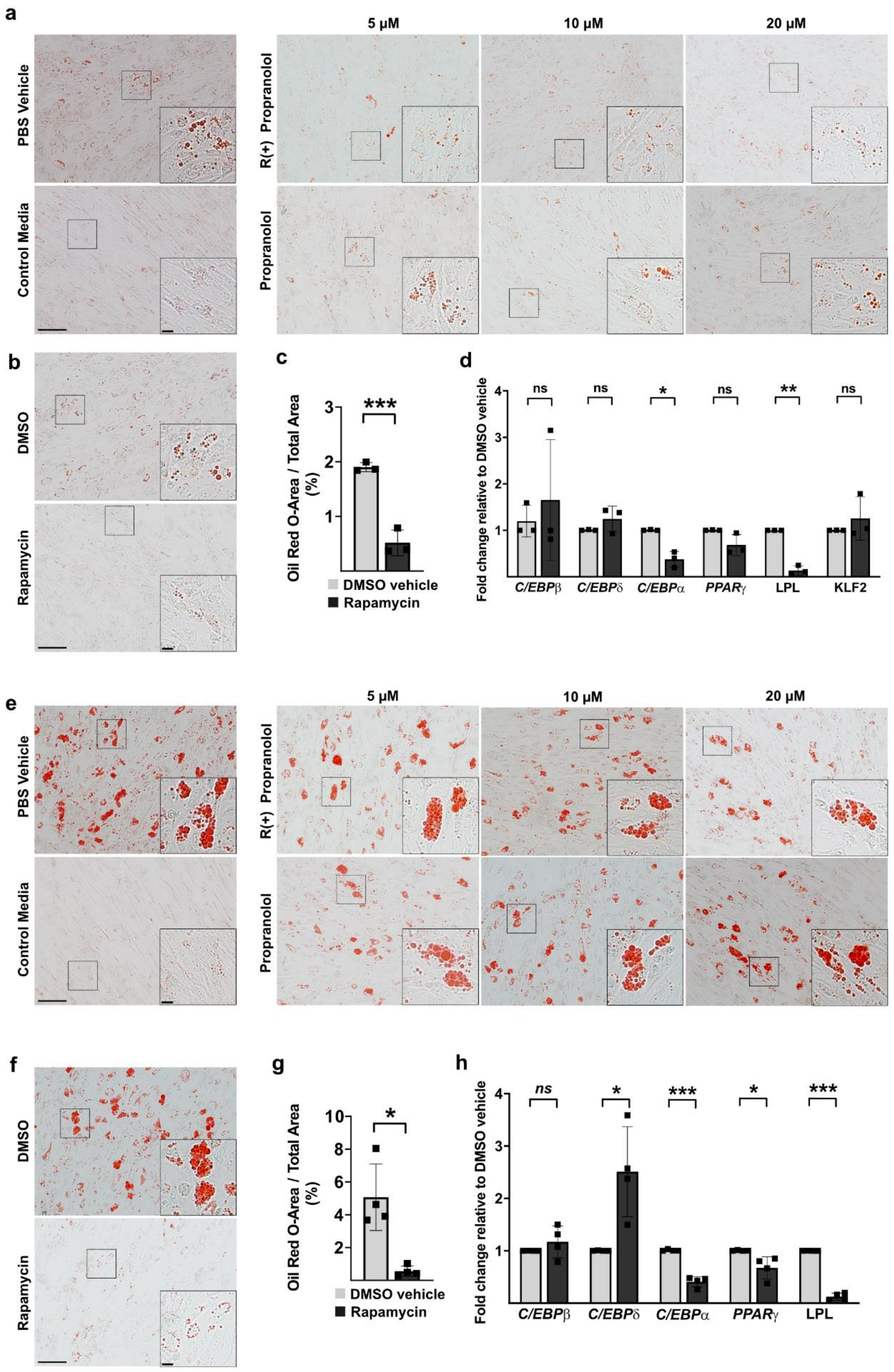

a
PBS Vehicle
Control Media
5 µM
10 µM
20 µM
R(+) Propranolol
Propranolol
b
DMSO
Rapamycin
c
Oil Red O-Area / Total Area (%)
***
DMSO vehicle
Rapamycin
d
Fold change relative to DMSO vehicle
ns
ns
*
ns
**
ns
C/EBPβ
C/EBPδ
C/EBPα
PPARγ
LPL
KLF2
e
PBS Vehicle
Control Media
5 µM
10 µM
20 µM
R(+) Propranolol
Propranolol
f
DMSO
Rapamycin
g
Oil Red O-Area / Total Area (%)
*
DMSO vehicle
Rapamycin
h
Fold change relative to DMSO vehicle
ns
*
***
*
***
C/EBPβ
C/EBPδ
C/EBPα
PPARγ
LPL

**Figure S2: R(+) propranolol and racemic propranolol reduce lipid accumulation in Hemangioma-derived stem cell (HemSC) induced for 4 days and 8 days of adipogenesis.** (a) Representative Oil Red O (ORO)-stained images for 0 – 20μM R(+) propranolol and racemic propranolol treated HemSC undergoing adipogenesis for 4 days. Scale bar = 100 μm. Magnified images from region of interest denoted by black dotted box. Scale bar = 15 μm. (b) ORO-stained images for 20 ng/ml rapamycin and DMSO-vehicle treated HemSC undergoing adipogenesis for 4 days. Scale bar = 100 μm. Magnified images from region of interest denoted by black dotted box. Scale bar = 15 μm. (c) Average ORO-stained area (% total) from (b). Data presented as mean ± SD, N = 3. ***P<0.001 compared to cells grown in DMSO vehicle in adipogenic media (AM). (d) Gene expression of CCAAT-enhancer-binding protein beta (C/EBPβ), C/EBPδ, C/EBPα, peroxisome proliferator-activated receptor gamma (PPARγ), lipoprotein lipase (LPL) and kruppel-like factor 2 (KLF2) determined by quantitative real time-polymerase chain reaction (RT-PCR). Data presented as mean ± SD, N = 3. *P<0.05, **P<0.01 compared to cells grown in DMSO vehicle in AM. (e) Representative ORO-stained images for 0 – 20uM R(+) propranolol and racemic propranolol treated HemSC undergoing adipogenesis for 8 days. Scale bar = 100 μm. Magnified images from region of interest denoted by black dotted box. Scale bar = 15 μm. (f) ORO-stained images for 20 ng/ml rapamycin and DMSO-vehicle treated HemSC undergoing adipogenesis for 8 days. Scale bar = 100 μm. Magnified images from region of interest denoted by black dotted box. Scale bar = 15 μm. (g) Average ORO-stained area (% total) from (f). Data presented as mean ± SD, N = 4. *P<0.05 compared to cells grown in DMSO vehicle in AM. (h) Gene expression of C/EBPβ, C/EBPδ, C/EBPα, PPARγ, LPL determined by qRT-PCR. Data presented as mean ± SD, N = 4. *P<0.05, **P<0.01 compared to cells grown in DMSO vehicle in AM.

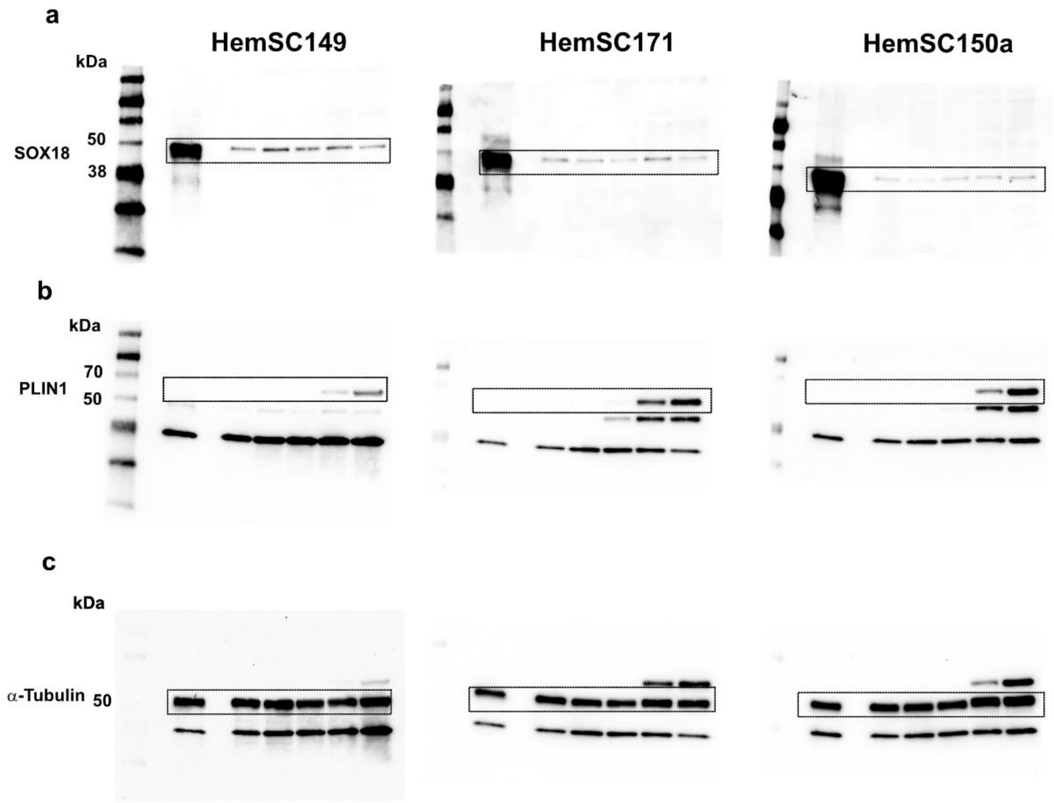


**Figure S3: Hemangioma-derived stem cells (HemSC) express basal levels of Sry-box transcription factor 18 (SOX18) protein during adipogenesis.** Full western blot images of HemSC (149, 171, 150a) induced for adipogenesis over 8-day time course and naive HemEC171. (a) SOX18 levels (b) Perilipin-1 (PLIN1) levels (c) α-tubulin levels.

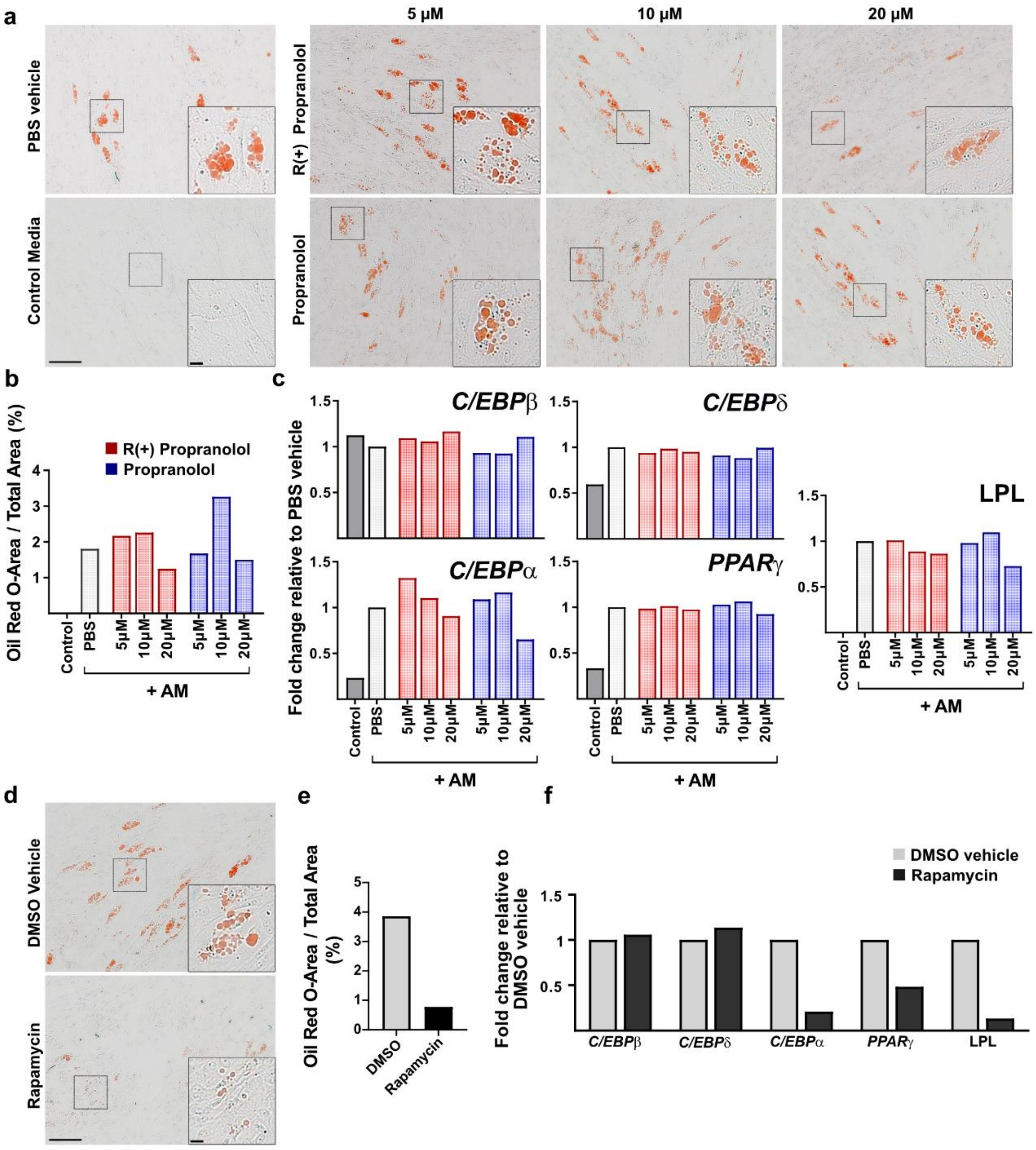


**Figure S4: R(+) propranolol and racemic propranolol effect on human white adipose tissue-derived mesenchymal stem cells (AMSC) induced for 8 days of adipogenesis.** (a) Representative Oil Red O (ORO)-stained images for 0–20μM R(+) propranolol and racemic propranolol treated AMSC undergoing adipogenesis for 8 days. Scale bar = 100 μm. Magnified images from region of interest denoted by black dotted box. Scale bar = 15 μm. (b) ORO-stained area (% total) from (b). N = 1. (c) Gene expression of CCAAT-enhancer-binding protein beta (C/EBPβ), C/EBPδ, C/EBPα, peroxisome proliferator-activated receptor gamma (PPARγ) and lipoprotein lipase (LPL) determined by quantitative real time-polymerase chain reaction (qRT-PCR). N = 1. AM = adipogenic media. (d) Representative ORO-stained images for 20 ng/ml rapamycin and DMSO vehicle-treated AMSC undergoing adipogenesis for 8 days. Scale bar = 100 μm. Magnified images from region of interest denoted by black dotted box. Scale bar = 15 μm. (e) ORO-stained area (% total) from (d). N=1. (f) Gene expression of C/EBPβ, C/EBPδ, C/EBPα, PPARγ, LPL determined by qRT-PCR. N=1.

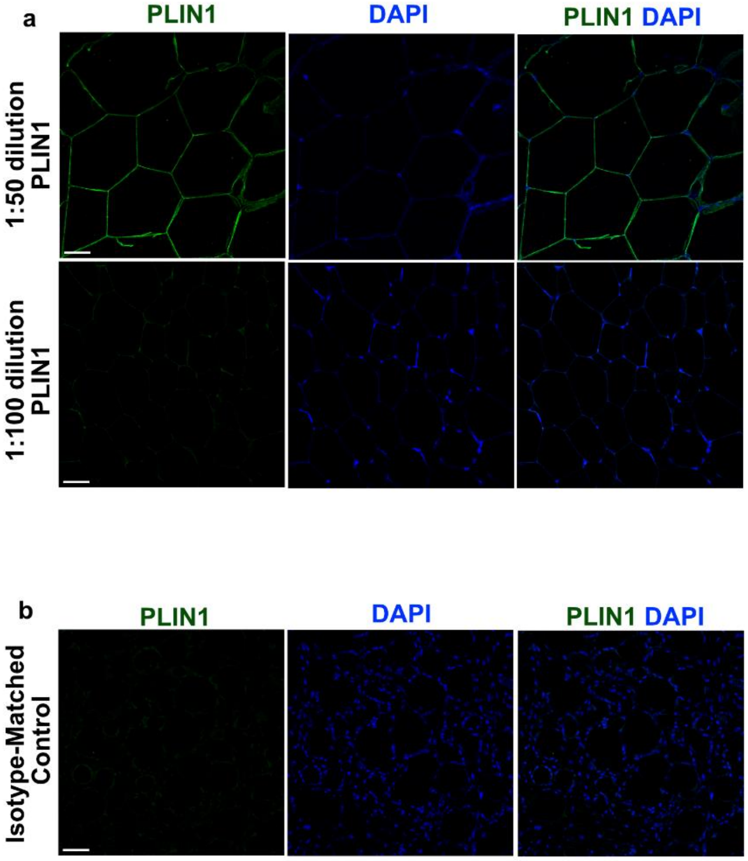


**Figure S5: Anti-human Perilipin-1 (PLIN1) antibody test.** (a) Anti-PLIN1 antibody titration immunohistochemistry (IHC) test on formalin-fixed paraffin-embedded (FFPE) sections from murine mammary fat from a FvB mouse strain at a dilution factor of 1:50 and 1:100 human PLIN1 antibody. Scale bar = 50 µm. (b) IHC on FFPE sections from an infantile hemangioma matrigel plug (untreated group) tissue with isotype-matched secondary antibody. Scale bar = 50 µm.

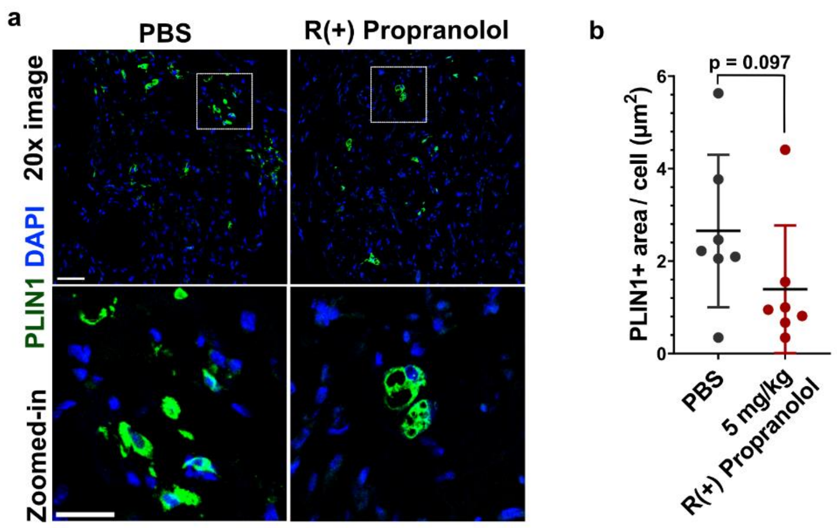


**Figure S6: 5 mg/kg R(+) propranolol decreases lipid accumulation in pre-clinical infantile hemangioma (IH) xenograft murine model.** Mice treated with 5 mg/kg R(+) propranolol versus phosphate-buffered saline (PBS) vehicle twice a day for 7 days. Hemangioma stem cell (HemSC) xenograft implants harvested for immunofluorescent staining. (a) Immunofluorescence of formalin-fixed paraffin-embedded sections stained with anti-human perilipin-1 (PLIN1). Top: Representative 20x images. Scale bar = 50 µm. Bottom: Magnified images at region of interest shown by white box. Scale bar = 25 µm. (b) Quantification of PLIN1-stained area per cell (µm$^2$) of implants in (a) using ImageJ. Data presented as mean ± SD, n = 7.

**SUPPLEMENTARY METHODS**

**Appendix S1**

**In vivo murine model for IH vasculogenesis**

The IH xenograft experiment was conducted by Seebauer et al. (2022). R(+) propranolol stock solution was prepared in double deionized $H_2O$ and diluted with phosphate-buffered saline accordingly. Hemangioma stem cells (HemSC) were grown in Endothelial Growth Media (EGM-2) supplemented with 10 µM R(+) propranolol, propranolol or PBS as a vehicle control for 24 hours before harvesting. HemSC were harvested and suspended in 200 µl Matrigel (Corning, USA) with 1 µg/ml erythropoietin (EPO; ProSpec, USA) and 1 µg/ml basic fibroblast growth factor (bFGF; ProSpec). A total of 3 x $10^6$ HemSC were introduced per implant. Each Matrigel/cell suspension was injected subcutaneously into the backs of 6-week-old male athymic nude mice (Massachusetts General Hospital, USA), totaling two implants per mouse. The mice were given 12.5 mg/kg propranolol, R(+) propranolol, or PBS as vehicle, twice each day via intraperitoneal injections. After the 7-day drug treatment, the mice were euthanized and implants removed for formalin fixation and paraffin embedment. The murine xenograft tissue slides were used for immunohistology by staining with anti-human perilipin-1 (PLIN1) to quantify lipid content.

**Western Blot**

Cells were lysed in a RIPA lysis and extraction buffer (Thermo Scientific, USA) supplemented with protease and phosphatase inhibitor cocktail (Cell Signalling Technology, USA). Equal protein amounts, determined by Bradford Assay (Bio-Rad, USA), were separated by SDS-PAGE and transferred to polyvinylidene difluoride membranes using Trans-Blot Turbo Transfer system (Bio-Rad, USA). Membranes were blocked with 5% nonfat milk for 1 hour at RT, incubated with primary antibodies overnight at 4$^{o}$C, washed thrice with tris-buffered saline-0.1% Tween-20 (TBST), incubated with secondary antibodies at RT for 1 hour, washed with TBST once for 15 minutes, five times with TBST for 5 minutes, and visualized using ECL detection reagent (Bio-Rad, USA).

**Antibodies used**

The antibodies used in this study are detailed here: mouse anti-human SOX18 (1:500; #SC-166025, Santa Cruz Biotechnology, USA), rabbit anti-human PLIN1 (1:1000, #3470, Cell Signaling Technology, USA), rabbit anti-human α-tubulin (1:5000, #2125, Cell Signaling Technology, USA), Goat anti-rabbit IgG, HRP-linked

Antibody (1:3000, 7074S, Cell Signaling Technology, USA), Horse anti-mouse IgG, HRP-linked Antibody (1:3000, 7076S, Cell Signaling Technology, USA).
